# Somatic DNA demethylation generates tissue-specific methylation states and impacts flowering time

**DOI:** 10.1101/2021.03.29.437569

**Authors:** Ben P. Williams, Lindsey A. Bechen, Deborah A. Pohlmann, Mary Gehring

## Abstract

Cytosine methylation is a reversible epigenetic modification to DNA. In plants, removal of cytosine methylation is accomplished by the four members of the DME family of 5-methylcytosine DNA glycosylases. Demethylation by DME is critical for seed development. Consequently, determining the function of the entire gene family in somatic tissues by mutant analysis has not been possible. Here, we bypassed the reproductive defects of *dme* mutants to create somatic quadruple homozygous mutants of the entire DME family. *dme; ros1; dml2; dml3* (*drdd*) leaves exhibit hypermethylated genomes compared to both wild-type plants and *rdd* triple mutants, indicating functional redundancy among all four demethylases. Targets of demethylation include regions co-targeted by RNA-directed DNA methylation and, surprisingly, CG gene body methylation, indicating dynamic methylation at these little-understood sites. Additionally, many tissue-specific methylation differences are absent in *drdd*, suggesting a role for active demethylation in generating divergent epigenetic states across wild-type tissues. Furthermore, *drdd* plants display a striking early flowering phenotype, which is associated with 5’ hypermethylation and transcriptional down-regulation of *FLOWERING LOCUS C*. Active DNA demethylation is therefore required for proper methylation patterning across somatic tissues and defines the epigenetic landscape of both intergenic and coding regions.

## Introduction

Phenotypic stability depends on epigenetic stability. Cytosine DNA methylation is both stably inherited and dynamic. In many plant species, DNA methylation is concentrated in transposable elements and other repetitive sequences, transcriptionally silencing these genomic regions. DNA methylation also accumulates in gene coding regions where it does not induce transcriptional silencing (Niederhuth et al., 2016; Bewick and Schmitz, 2017). Methylation patterns can be highly conserved over evolutionary time but also dynamic in specific developmental contexts, such as during reproduction (Gehring, 2019). These patterns are a result of methylation activity and demethylation activity, the coordination of which is required to maintain transgenerational epigenetic and phenotypic stability (Williams and Gehring, 2017). Methylation is established by *de novo* DNA methyltransferases, which are guided to their targets by small RNAs in a process termed RNA-directed DNA methylation (RdDM) (Matzke and Mosher, 2014). Symmetric DNA methylation is maintained by maintenance methyltransferase enzymes that copy patterns of DNA methylation after DNA replication. Loss of DNA methylation can occur passively, when DNA methylation is not maintained after DNA replication, or by active removal by DNA demethylation pathways. Active DNA demethylation is initiated by the activity of a family of bifunctional HhH-GPD DNA glycosylases/lyases (Gehring et al., 2006; Morales-Ruiz et al., 2006; Penterman et al., 2007; Ortega-Galisteo et al., 2008; Agius et al., 2006). In *Arabidopsis thaliana*, there are four gene family members: *DME, ROS1, DML2*, and *DML3* (Choi et al., 2002; Gong et al., 2002; Penterman et al., 2007; Ortega-Galisteo et al., 2008). These enzymes remove methylated cytosines through base excision repair (Roldán-Arjona et al., 2019). *DME* and *ROS1* genes are present in plants from algae onwards (Pei et al., 2019). The distribution of *DML2* and *DML3* is more limited, with *DML3* present in monocots and dicots and *DML2* present only in a subset of dicots (Pei et al., 2019).

Molecular and morphological phenotypes have been described for plants mutant in one or more of the 5-mC DNA glycosylases in Arabidopsis and rice. Arabidopsis plants heterozygous for *dme* mutations have a visibly striking phenotype: 50% seed abortion (Choi et al., 2002). Seeds that inherit a mutant *dme* allele from the mother abort after several days of development and thus homozygous *dme* mutants have only rarely been recovered. In some Arabidopsis accessions, *dme* is also not fully transmitted through the male parent (Schoft et al., 2011). DME is active in the polar nuclei and the central cell (the female gamete that is the progenitor of the endosperm) before fertilization and is required to establish gene imprinting in the endosperm after fertilization (Choi et al., 2002; Gehring et al., 2006). In the central cell, DME demethylates specific loci (Park et al., 2016) and this hypomethylated state is transmitted to the endosperm after fertilization, such that the maternally-inherited endosperm genome is hypomethylated compared to the paternally-inherited endosperm genome (Gehring et al., 2006, 2009; Hsieh et al., 2009; Ibarra et al., 2012). DME-dependent endosperm hypomethylated sites are enriched for fragments of transposable elements that reside near genes (Gehring et al., 2009). DME is also active in the pollen vegetative cell and similar targets are hypomethylated in a DME-dependent manner in both the vegetative cell and endosperm (Ibarra et al., 2012; Calarco et al., 2012). Like Arabidopsis, rice central cells and vegetative cells are hypomethylated (Park et al., 2016; Kim et al., 2019). Mutations in a rice *DME* homolog (termed *ROS1a*) have a similar phenotype as Arabidopsis *DME* mutants – maternal null alleles disrupt endosperm development and mutant maternal or paternal alleles are only rarely transmitted to progeny (Ono et al., 2012). *ROS1a* is also responsible for DNA demethylation in the vegetative cell (Kim et al., 2019) and likely also in the central cell. Although Arabidopsis DME is mostly highly expressed in reproductive tissues, expression is also detected in vegetative tissues (Mathieu et al., 2007; Park et al., 2017; Schumann et al., 2019). The function of DME outside of reproductive tissues, if any, is largely unknown.

Arabidopsis plants with mutations in any of the other three DNA glycosylases lack visibly dramatic phenotypes under standard growth conditions and all null mutants are viable singly or in combination (Gong et al., 2002; Penterman et al., 2007; Ortega-Galisteo et al., 2008). However, some phenotypes have been observed. Stomatal precursor-cell density is increased in *ros1* leaves due to hypermethylation and transcriptional silencing of a negative regulator of precursor cells (Yamamuro et al., 2014). Additionally, *ros1* and *ros1*; *dml2*; *dml3* (*rdd*) triple mutants exhibit impaired tracheary element differentiation, resulting in a high frequency of protoxylem discontinuities (Lin et al., 2020). Plants with mutations in *ros1* and *rdd* exhibit enhanced susceptibility to bacterial and fungal pathogens (Yu et al., 2013; Le et al., 2014; López Sánchez et al., 2016), which is associated with decreased expression of biotic stress genes and, in a handful of examined genes, promoter hypermethylation at TE sequences (Le et al., 2014; Halter et al., 2021). Genome-wide profiling of DNA methylation in *ros1* plants and *rdd* plants indicates DNA hypermethylation at hundreds or thousands of intergenic regions and transposable elements that are also targeted by RdDM and are enriched near genes (Penterman et al., 2007; Lister et al., 2008; Tang et al., 2016). Most of the DNA methylation changes in *rdd* are not obviously associated with changes in gene transcription (Penterman et al., 2007; Lister et al., 2008). Thus, it is thought that one function of the DNA demethylation in vegetative tissues is to ‘clean-up’ after a robust DNA methylation system to keep genes free from methylation.

Although genetic approaches have proved fruitful in understanding the function of 5-mC DNA glycosylases, the inability to generate homozygous mutant *dme* plants has stymied full understanding of this gene family. Recently, RNAi was used to knockdown DME expression in vegetative tissues of the *rdd* triple mutant (Schumann et al., 2019). These plants are even more susceptible to a fungal pathogen and additional hypermethylation was shown at a few loci. These data suggest that vegetative tissue of *rdd* plants retain some demethylation activity from *DME*. To identify the full extent of demethylation activity, we created plants where *dme* was complemented only in the central cell, allowing us to examine methylation and transcription on a genome-wide scale in vegetative tissues null for all four 5-methylcytosine DNA glycosylases.

## Results

### Isolating quadruple demethylase mutants

In order to fully understand the role of the DME family of 5-methylcytosine DNA glycosylases in Arabidopsis, we sought to isolate homozygous mutations in all four orthologues in this family – *DME, ROS1, DML2* and *DML3*. To bypass the *dme* seed abortion phenotype, we created a transgene in which the genomic coding sequence of DME was expressed under the central cell-specific promoter of *AGL61* (Fig. 1A) (Steffen et al., 2008). This transgene was transformed into the previously isolated *rdd* triple mutant (Penterman et al., 2007). Transgenic *rdd* plants were pollinated by heterozygous *dme* mutants to create heterozygous quadruple mutants (Fig. 1B). These heterozygotes were self-fertilized and seed abortion rates were quantified to test the ability of the *pAGL61:DME* transgene to complement the *dme* seed abortion phenotype. Whereas non-transgenic plants harboring the *dme* mutation exhibit 50% seed abortion, multiple transgenic lines expressing *pAGL61:DME* exhibited minimal seed abortion, indistinguishable from *rdd* plants. Expression of *DME* in the central cell before fertilization is therefore sufficient to rescue the post-fertilization seed abortion phenotype. After self-fertilizing the heterozygous quadruple mutant, the following four genotypes were isolated over two subsequent generations: homozygous quadruple mutants (hereafter termed *drdd*), homozygous *rdd* mutants, homozygous *dme* mutants, and homozygous wild-type segregants (hereafter termed WT, and to serve as genetically closely related wild-type controls for subsequent experiments). All four of these genotypes were determined to be homozygous for a single transgene insertion, and displayed seed abortion rates of <2%, similar to non-transgenic *rdd* (Fig. 1C).

**Figure 1.**
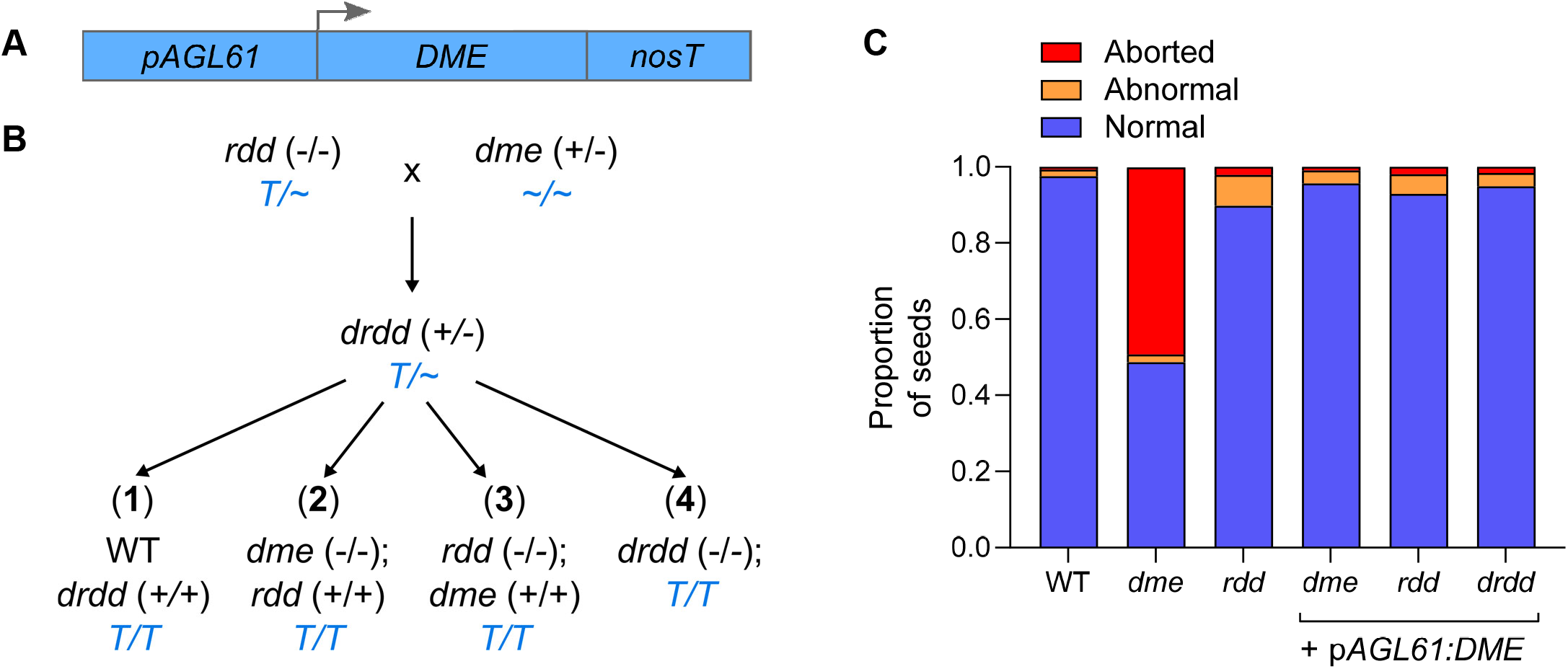
Generation of homozygous *dme* and quadruple *drdd* mutants. A) Schematic showing the construct designed to express *DME* specifically in central cells using the *AGL61* promoter. B) Schematic showing the segregation of mutant genotypes. Transgenic *rdd* plants were pollinated by heterozygous (+/-) *dme* to generate heterozygous *drdd* F_1_ progeny. WT, *dme, rdd* and *drdd* homozygotes were isolated from two subsequent generations. T and ∼ refer to the prescence and absence of the *pAGL61:DME* transgene respectively. C) Proportion of aborted seeds in non-transgenic WT, *dme, rdd* plants, as well as *dme, rdd* and drdd plants homozygous for the *pAGL61:DME* transgene. 250-300 seeds were evaluated for each genotype.

### DME is required for full demethylation in somatic tissue

To fully understand the function of *DME* outside of the central cell, and the combined activity of all members of the DME family, paired DNA and RNA samples were collected from leaf tissue of WT, *dme, rdd* and *drdd* plants. Individual leaves from four separate biological replicate plants were sliced along the midvein, and DNA and RNA were extracted from one half each, so that DNA methylation and gene expression could be analyzed from the same leaf.

Whole genome bisulfite sequencing (BS-seq) was performed on DNA samples. In order to improve read depth for subsequent analyses, bisulfite-sequencing reads were combined for pairs of replicates to create two high-depth replicates for each genotype. Previously, the *rdd* triple mutant was constructed by back-crossing *ros1* and *dml2* T-DNA insertions originally from a Ws-2 background into a Col-0 background (Penterman et al., 2007). We therefore first analyzed the zygosity of Ws-2 SNPs throughout the sequenced WT, *dme, rdd*, and *drdd* methylomes to determine the presence of chromosomal regions originating from Ws-2. Consistent with previous reports (Penterman et al., 2007), we observed regions of Ws-2 homozygosity surrounding the *ros1* and *dml2* T-DNA insertions on chromosomes 2 and 3 respectively, both in *rdd* and *drdd* (Supp. Fig. 1). These regions were discarded in all methylation analyses, as strain-specific methylation differences between Col-0 and Ws-2 could contribute to false positives in identifying differentially methylated regions.

In order to assess the extent of methylation differences in *dme, rdd*, and *drdd*, differentially methylated dinucleotides (CG context) or individual cytosines (CH context) were identified between WT and each mutant genotype, selecting only for CGs/Cs that were differentially methylated in both mutant replicates (see methods). Methylation differences were restricted to a minimum percentage difference of 35% for CG, 20% for CHG and 15% for CHH. Differentially methylated CGs/Cs (DMCs) were then aggregated into windows, which were assigned a differential methylation score based on the quantity and proportion of DMCs (Williams and Gehring, 2017). As this approached required DMCs to have adequate read depth in both replicates in order to be counted, we estimate that our analysis has few false positives, but as a consequence likely offers an under-estimate of the true number of methylation differences across the mutant genomes.

Using this approach, we identified few differentially methylated regions (DMRs) between WT and *dme* mutants (Fig. 2B). This suggests that DME does not have a significant number of target loci that are independent of the other three demethylases. We observed approximately 2,000 regions with increased methylation in *rdd* compared to WT. The quadruple mutant *drdd* exhibited even greater methylation differences compared to WT, with approximately 1,000 loci more than *rdd*, including almost twice the number of regions with differential non-CG methylation. This suggests that DME does function in somatic tissue, but acts redundantly with ROS1, DML2, and DML3. The set of loci hypermethylated in *drdd* compared to wild-type therefore represents a set of genomic targets for the entire DME family, which we hereafter refer to as “DRDD target loci” (Supp. Dataset 1). To better understand the extent of DME activity at DRDD target loci, we quantified methylation levels at these regions in all four genotypes studied. Single mutant *dme* replicates had methylation levels similar to WT, suggesting that DME functions redundantly with the other demethylases in removing methylation at these targets (Fig. 2C). Whereas *rdd* triple mutants showed hypermethylation in all three contexts at DRDD target loci, this hypermethylation was much more pronounced in *drdd* (Fig. 2C). At the single-cytosine level, approximately 70% of CGs within DRDD target loci exhibited increased methylation in *drdd* compared to *rdd*, as well as >50% of CHG and CHH cytosines, even though many of these methylation changes were smaller than the cutoffs of our prior analysis. These data suggest that in *rdd* plants, DME still actively removes methylation from the majority of DRDD target loci.

**Figure 2.**
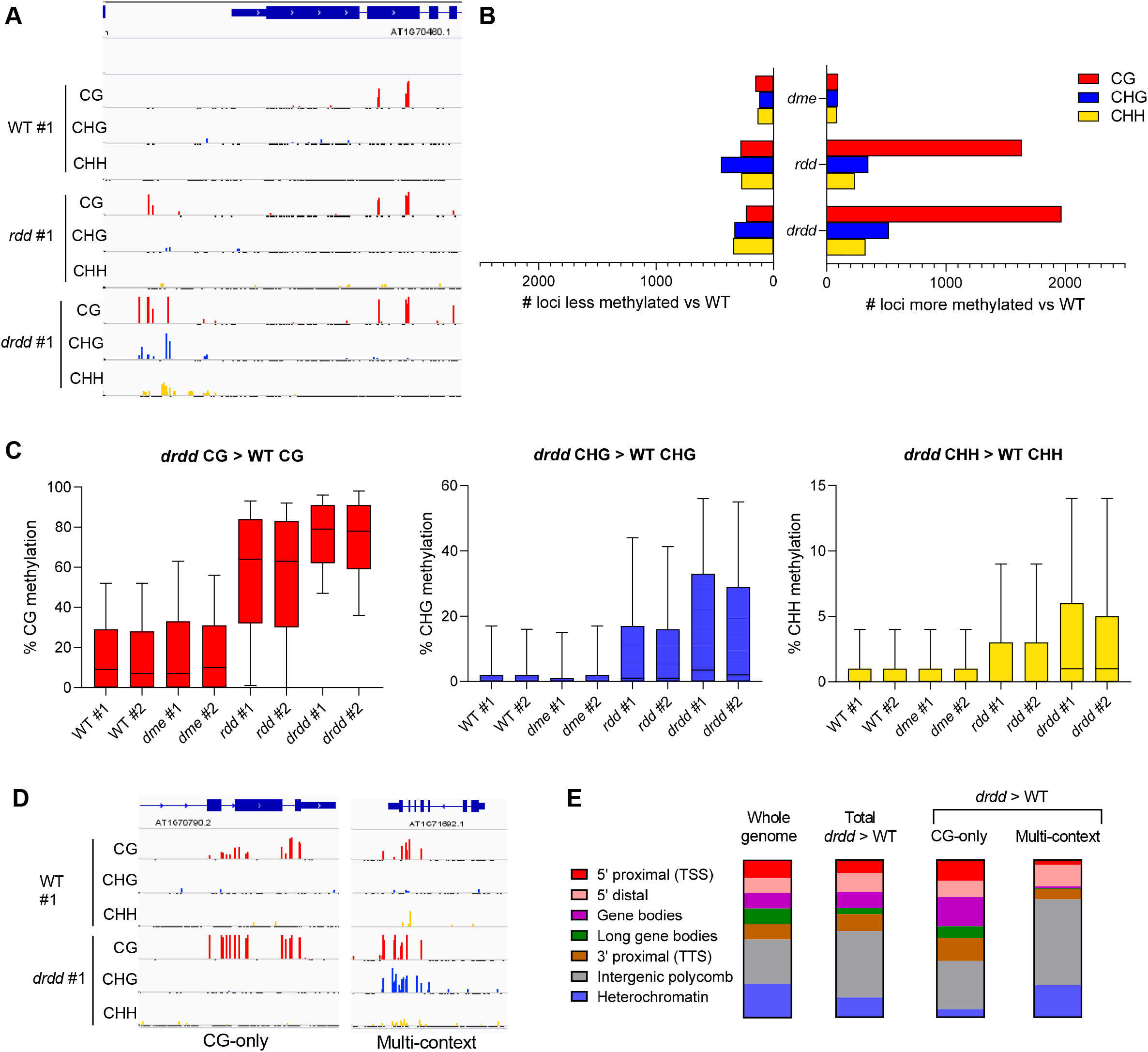
Exacerbated DNA hypermethylation in *drdd* mutants. Genome browser snapshot showing typical locus hypermethylated in *drdd* compared to WT and *rdd*. B) Number of differentially methylated regions in *dme, rdd* and *drdd* compared to WT. C) Methylation levels of WT, *dme, rdd* and *drdd* at loci hypermethylated in *drdd* compared to WT. Boxes denote the interquartile range, whiskers denote 10th and 90th percentiles. D) Genome browser snapshots showing typical examples of a locus hypermethylated specifically in the CG context (CG-only) and a locus hypermethylated in all sequence contexts (multi-context). E) The distribution of loci hypermethylated in *drdd* compared to WT across different genomic chromatin states.

We next sought to better understand the genomic context of DRDD target loci, as well as their characteristic DNA methylation patterns. To do this, we identified the chromatin state of each DRDD target using a published dataset of chromatin states that incorporates information from 16 characteristics and histone modifications (Sequeira-Mendes et al., 2014). The overall set of DRDD targets was dispersed over a distribution of chromatin states highly similar to the entire Arabidopsis genome (Fig. 2E). DRDD target loci could be classified in either of two groups: “CG-only” loci, which lack non-CG methylation in both WT and *drdd*, and exhibit increased CG methylation in *drdd*, and “multi-context” loci, which exhibit hypermethylation in CG, CHG and CHH contexts in *drdd* (Fig. 2D). While equally common, these two types of target loci occupied different chromatin states. CG-only DMRs were more commonly associated with gene bodies and the immediately adjacent 5’ and 3’ regions, whereas multi-context loci where more common in distal 5’ regions and other intergenic DNA associated with the repressive histone mark H3K27me3 (Fig. 2E). These differences in chromatin state distribution likely reflect the well-characterized differences between the distributions of gene body methylation (gbM), which is typically only on CG-dinucleotides, and the activities of the RdDM and CMT2 pathways, which are active at repetitive DNA and intergenic sequences (Stroud et al., 2014; Bewick and Schmitz, 2017). We note that these CG-only DMRs are unlikely to represent the trivial differences in CG methylation that can spontaneously arise between generations, often termed spontaneous epimutations (Johannes and Schmitz, 2019). Our approach to identifying DMRs was designed to avoid identifying large numbers of epimutations, as evidenced by the small number of significant CG methylation differences identified between WT and *dme* (<100) which shared the same grand-parental heritage as WT and *drdd*. These CG-only target loci therefore represent loci specifically targeted for demethylation in leaf tissues, raising the possibility that CG-gene body methylation may play a functional role in some cell or tissue types.

### Demethylation by DRDD establishes tissue-specific methylation states

DRDD target loci exhibit low but often non-zero levels of CG methylation in wild-type leaves which are by definition hypermethylated in *drdd*. This suggests that both methylation and demethylation processes occur at these sites in wild-type rosette leaves (Figs. 2C & 3A). The low level of CG methylation at DRDD target loci in wild-type leaves is in contrast to most CG sites in the genome, which are either not methylated or fully methylated (Fig. 3A). We hypothesized that the balance between methylation and demethylation occurring at DRDD target loci could be altered in different tissues or over developmental time. To test this, we generated whole genome bisulfite sequencing data for tissues from individual wild-type Col-0 plants, including rosette leaves (of the identical developmental stage as our prior leaf samples), cauline leaves, closed flower buds, and mature green embryos. Cauline leaves and flower buds were collected on the same day. Two replicates were sequenced, with each set of tissues collected from one individual plant.

**Figure 3.**
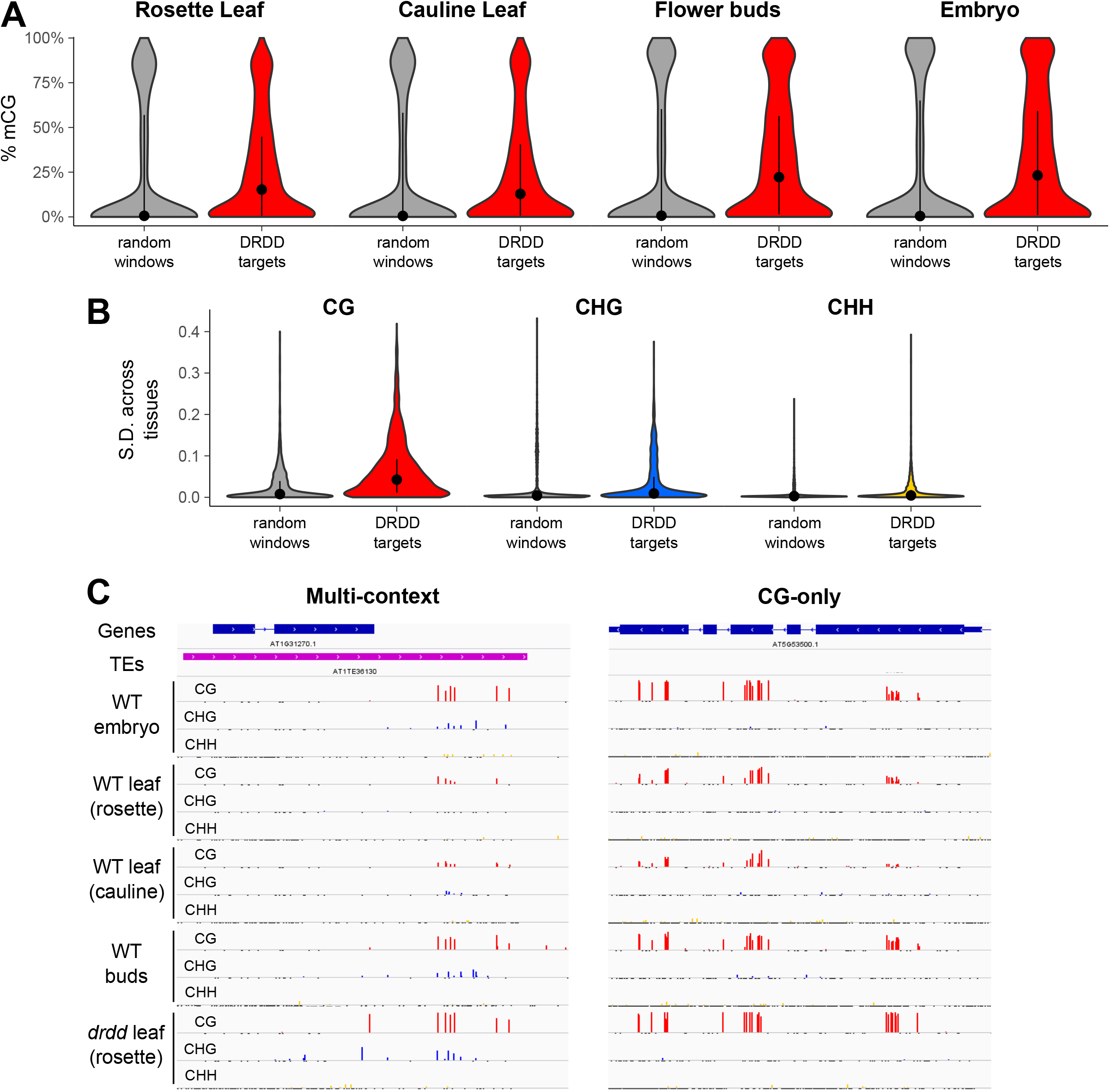
Demethylation by DRDD establishes tissue-specific methylation differences. A) CG methylation levels of four different WT tissues at 2,601 DRDD target loci, compared with 2,601 randomly-selected genomic windows. B) Standard deviation of DNA methylation levels across tissues at DRDD target loci compared to random windows. C) Genome browser snapshots depicting typical examples of multicontext and CG-only DRDD targets in different WT tissues.

We identified hundreds of regions differentially methylated at CG dinucleotides among tissues, and thousands of regions differentially methylated in CHG or CHH contexts (Supp. Fig. 2). Embryos exhibited approximately 4,000 regions with increased CHH methylation when compared to rosette or cauline leaves, suggesting increased activity of the RdDM or CMT2 pathways at this developmental stage, consistent with previous reports documenting embryo hypermethylation (Narsai et al., 2017; Kawakatsu et al., 2017; Bouyer et al., 2017). In addition, we observed increased CHG methylation at over 10,000 regions in embryos or flower buds compared to both rosette and cauline. CHH and CHG methylation sites typically exhibit partial methylation, with only a fraction of sites methylated in whole-tissue bisulfite sequencing data, suggesting cellular heterogeneity. It is therefore possible that differences in the cell type composition of different tissues could cause large differences in the levels of non-CG methylation observed in whole tissues. Rosette and cauline leaves exhibited highly similar methylation profiles in our dataset, suggesting that cell-type composition is predictor of methylation patterns.

To evaluate the relationship between tissue-specific methylation differences and DRDD, we calculated the methylation levels of 2,601 DRDD target loci in each tissue of our dataset, as well as a set of 2,601 randomly selected 200 bp windows that fell within the statistical boundaries necessary to be identified as a hypermethylated region (hyperMR). DRDD target loci exhibited increased methylation in flower buds and embryos compared to the leaf samples, with the largest differences in CG methylation at CG-only DRDD target loci (Fig. 3A, Supp. Fig. 3). Tissue-type differences were not observed in the randomly-selected windows, suggesting the observed differences are specific to DRDD target loci. We also calculated the standard deviation in DNA methylation among tissues at hyperMRs and control regions. The standard deviation of methylation levels between tissues was much higher at DRDD target loci than at randomly selected windows (Fig. 3B), suggesting that DRDD targets overlap with loci that exhibit developmentally dynamic epigenetic states. Lastly, >50% of regions identified to be CG hypomethylated in (either rosette or cauline) leaves compared to buds or embryos coincided with DRDD target loci (Supp. Fig. 2). Thus, hundreds of loci exhibited high methylation levels in WT buds and embryos but reduced methylation in cauline and rosette leaves, as well as high methylation levels in *drdd* rosette leaves (Fig. 3C). We observed this DRDD-dependent demethylation in WT leaves at both CG-only and multi-context DRDD targets (Supp. Fig. 3). To assess whether the activity of the RdDM pathway might also play a role in establishing these tissue specific-differences in methylation state, we sequenced the methylome of the same four tissues from *rdr2* mutants. *RDR2* is involved in biogenesis of the small RNAs that direct DNA methylation. Multi-context DRDD targets were not more methylated in *rdr2* flower buds and embryos than rosette and cauline leaves, suggesting that an interplay between RdDM and DRDD-mediated demethylation is responsible for establishing methylation differences across tissues at these sites (Supp. Fig. 3). In contrast, the methylation profile of CG-only DMRs across tissues was largely unchanged in *rdr2* compared to WT, suggesting that active demethylation by DRDD establishes CG methylation differences independently of RdDM. Together, these data suggest that active demethylation by DRDD is responsible for removing methylation at target loci specifically in leaf tissues. Our study has therefore uncovered previously unidentified developmental regulation of epigenetic states.

### Demethylation by DRDD is associated with target gene regulation

DNA methylation has variable impacts on gene expression. While many genes are not affected by changes to DNA methylation at proximal sequences, a subset of genes are strongly impacted by DNA methylation, typically by transcriptional silencing and/or heterochromatin formation (Lister et al., 2008), although occasionally by promoting increased expression (Williams et al., 2015; Pignatta et al., 2018).

To identify which genes might be impacted by the activity of DRDD demethylases, we performed RNA-seq on WT, *dme, rdd*, and *drdd* tissue isolated from the same leaf samples used for methylation profiling. WT and *dme* transcriptomes were almost identical (Fig. 4A), consistent with the minimal differences observed in methylation patterns between the two genotypes. We identified 168 downregulated and 174 upregulated genes in *rdd* compared to WT, and 170 downregulated and 252 upregulated genes in *drdd* compared to WT. Overall, the *drdd* transcriptome encompassed a larger dynamic range of expression changes relative to WT, suggesting that the most extreme expression changes were exacerbated in *drdd* compared to *rdd* (Fig. 4A, Supp. Fig. 4, Supp. Dataset 2).

**Figure 4.**
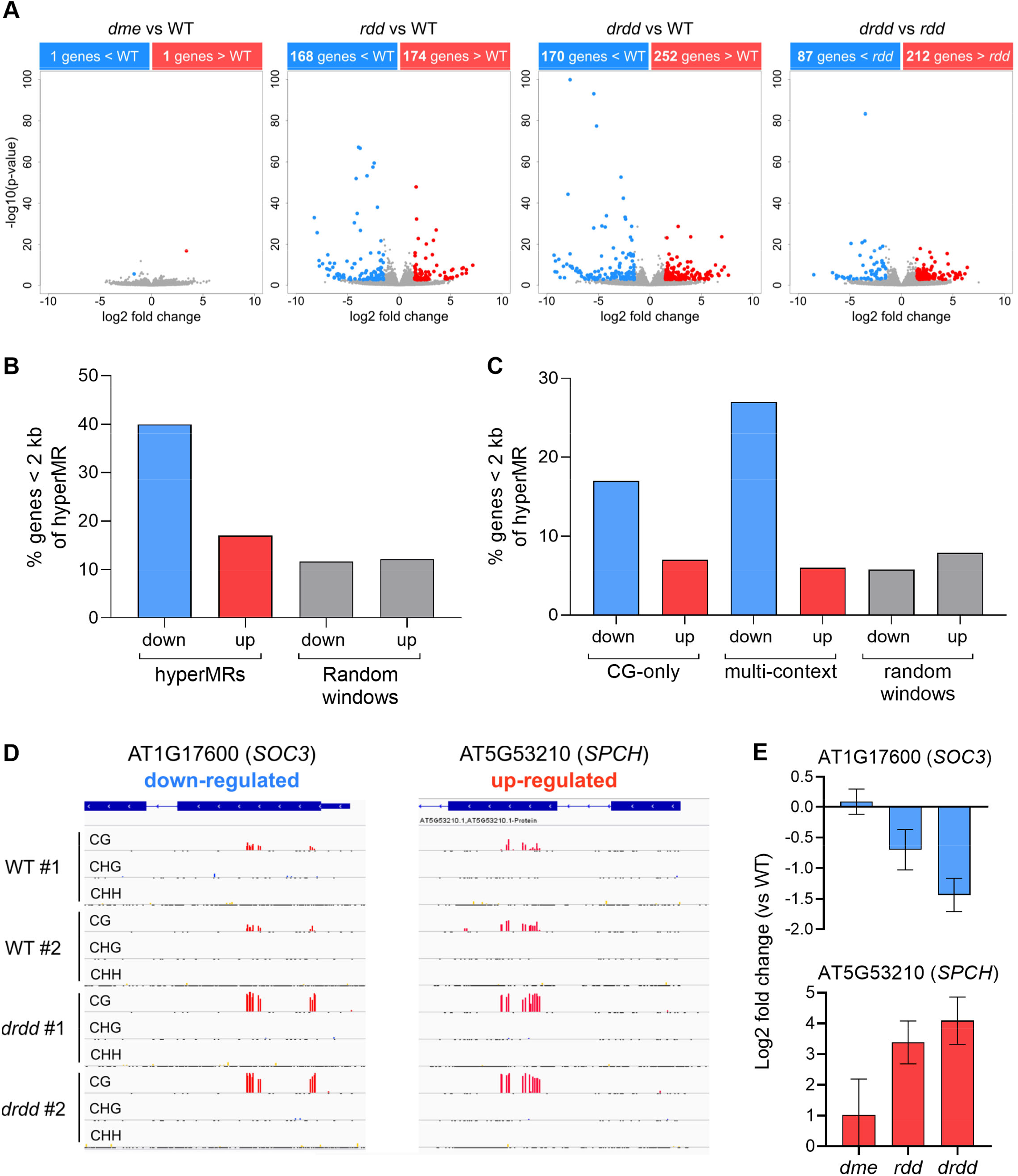
Demethylation by DRDD directly impacts expression of a subset of target genes. A) Volcano plots showing differentially expressed genes between *dme, rdd* and *drdd* compared to WT, as well as *drdd* compared to *rdd*. B) Percentage of differentially expressed genes within 2kb proximity to regions hypermethylated in *drdd* compared to WT. Proximity between differentially expressed genes and randomly selected genomic windows is shown as a negative control. C) Percentage of differentially expressed genes within 2kb proximity of CG-only or multi-context DRDD target loci. D) Genome browser snapshots showing two examples of CG-only DRDD target loci that are also differentially expressed. E) Gene expression differences of example CG-only target genes in *dme, rdd* and *drdd* compared to WT. Error bars represent standard error between biological replicates.

To examine whether these expression changes were linked to DNA methylation differences between *drdd* and WT, we calculated the distance between up and down-regulated genes and the closest DMR. Regions of increased DNA methylation (hyperMRs) were frequently adjacent to down-regulated genes, with 40% of down-regulated genes residing within 2 kb of a hyperMR (Fig. 4B). This association is consistent with previous studies of *ros1* and *rdd* mutants, which suggested that a small subset of methylation gains in these mutants caused transcriptional silencing of proximal genes (Lister et al., 2008; Yamamuro et al., 2014; Halter et al., 2021). We also observed proximity between hyperMRs and some up-regulated genes, with 14 % of up-regulated genes residing <2 kb of a hyperMR. However, this number is similar to that expected by chance, as determined by calculating the distance between hyperMRs and control groups of randomly selected 200 bp windows (Fig. 4B).

The proximity between a subset of hyperMRs and down-regulated genes was detectable for both multi-context and CG-only hyperMRs (Fig. 4C). An association between down-regulated genes and multi-context hyperMRs was expected, as methylation in multiple sequence contexts is a hallmark of the RdDM pathway, which typically functions in transcriptional silencing. The association of CG-only DMRs with decreased expression in *drdd* was more surprising, as CG gene body methylation has mostly been associated with minimal impacts on gene regulation in plants (Bewick and Schmitz, 2017; Picard and Gehring, 2017). For example, the locus encoding the defense and cell death regulator *SOC3* is down-regulated in *drdd* mutants, and exhibits clear hypermethylation of CG dinucleotides within the first exon (Fig. 4D-E, Supp. Fig. 5). Conversely, we also observed that increased *SPEECHLESS* expression in *drdd* was associated with increased CG gene body methylation (Fig. 4D-E). DRDD may actively remove methylation from these coding regions to facilitate higher or lower expression, or the altered transcriptional output of these genes may affect a different epigenetic state (Fig. 4D-E, Supp. Fig. 5). The demethylating activities of DRDD enzymes within coding regions as well as upstream sequences may be required to protect transcriptional activity from the over-reach of methylation pathways.

To better understand if the gene expression differences between WT and *drdd* mutants are associated with any particular biological functions, we performed GO term analysis (Supp. Fig. 4). Genes with decreased expression in *drdd* were significantly associated with one GO term, immune response, which was also associated with genes that increased in expression (Supp. Fig. 4). Several defense-response genes down-regulated in *drdd* were previously identified as targets of ROS1 and are in transposon-rich rapidly evolving regions of chromosomes, which tend to be frequently targeted by the RdDM pathway (Le et al., 2014; Halter et al., 2021). We observed greater increases in the expression of epidermal development genes in *drdd* than in *rdd* (Supp Fig. 4). GO terms for regulation of hormone levels, response to osmotic stress and response to cold were also significantly enriched among up-regulated genes (Supp. Fig. 4).

### DRDD enzymes remove methylation from the FLC locus and are required to delay flowering

During long-day (16 h) growth conditions, we observed that *drdd* mutants were early flowering compared to WT, *dme*, and *rdd* plants, typically transitioning to flowering after establishing 8-9 true leaves (Fig. 5A, Supp. Fig. 6). Under short day (8 h) growth conditions, we observed an early flowering phenotype both in *drdd* and *rdd*, which has not been noted in previous studies (Fig. 5B, Supp. Fig. 6). We therefore hypothesized that active DNA demethylation in Arabidopsis might regulate flowering time, perhaps by altering the epigenetic state at genes important for regulating the transition to flowering. In our RNA-seq dataset, the flowering time regulator *FLC* had significantly reduced expression in comparisons of WT vs *drdd* and *rdd* vs *drdd*. In fact, *FLC* transcripts were not detected in *drdd* mutants (Fig. 5C). We then sought to verify the expression levels of FLC using quantitative real-time PCR. Although we observed a high variance in *FLC* transcript abundance between biological replicates, *FLC* expression was clearly reduced in *drdd* mutants compared to WT, with >20-fold lower median expression value (Fig. 5D). Complex epigenetic regulation of the *FLC* locus by histone modification pathways has been well-described (Wu et al., 2020), but a clear connection to *cis* DNA methylation has not previously been reported.

To assess the DNA methylation state of the *FLC* locus, we analyzed the DNA methylation profiles of WT, *dme, rdd* and *drdd* plants in our BS-seq data. We observed minimal DNA methylation in the transcribed region or 3’ region of *FLC*, but saw evidence of DNA methylation in all 3 sequence contexts in a region 1,121-2,139 bp 5’ of the transcriptional start site of the *FLC* coding region. To better define the methylation patterns of this region with greater depth, we performed bisulfite PCR (BS-PCR) on three separate fragments. For two of these fragments, we generated data for both strands so that we could assay asymmetric CHH methylation accurately. In all BS-PCRs, we observed no methylation in WT or *dme* plants, consistent with the observation that *dme* mutants do not exhibit a flowering time phenotype (Fig. 5B, E). Across the three regions, we observed varying levels of methylation in all three sequence contexts in both *rdd* and *drdd. drdd* mutants exhibited the highest methylation at the more 5’-distal sequences, whereas *rdd* exhibited higher methylation than *drdd* between 1,121-1,434 bp upstream of the *FLC* coding region. Thus, DRDD enzymes target the region 5’ of *FLC* for demethylation. Hypermethylation of this region is correlated with reduced *FLC* expression, as well as reduced flowering time. As this region is highly methylated in all three sequence contexts, we hypothesize that it is targeted by RNA-directed DNA methylation. Indeed, 23-24 nucleotide small RNAs homologous to this precise region are present within multiple WT small RNA datasets (Lee et al., 2012; Jeong et al., 2013; Erdmann et al., 2017), which could direct RdDM. Thus, *FLC* is an exemplar of a locus that lacks DNA methylation in wild-type leaves due to active DNA demethylation, rather than lack of methylation targeting.

**Figure 5.**
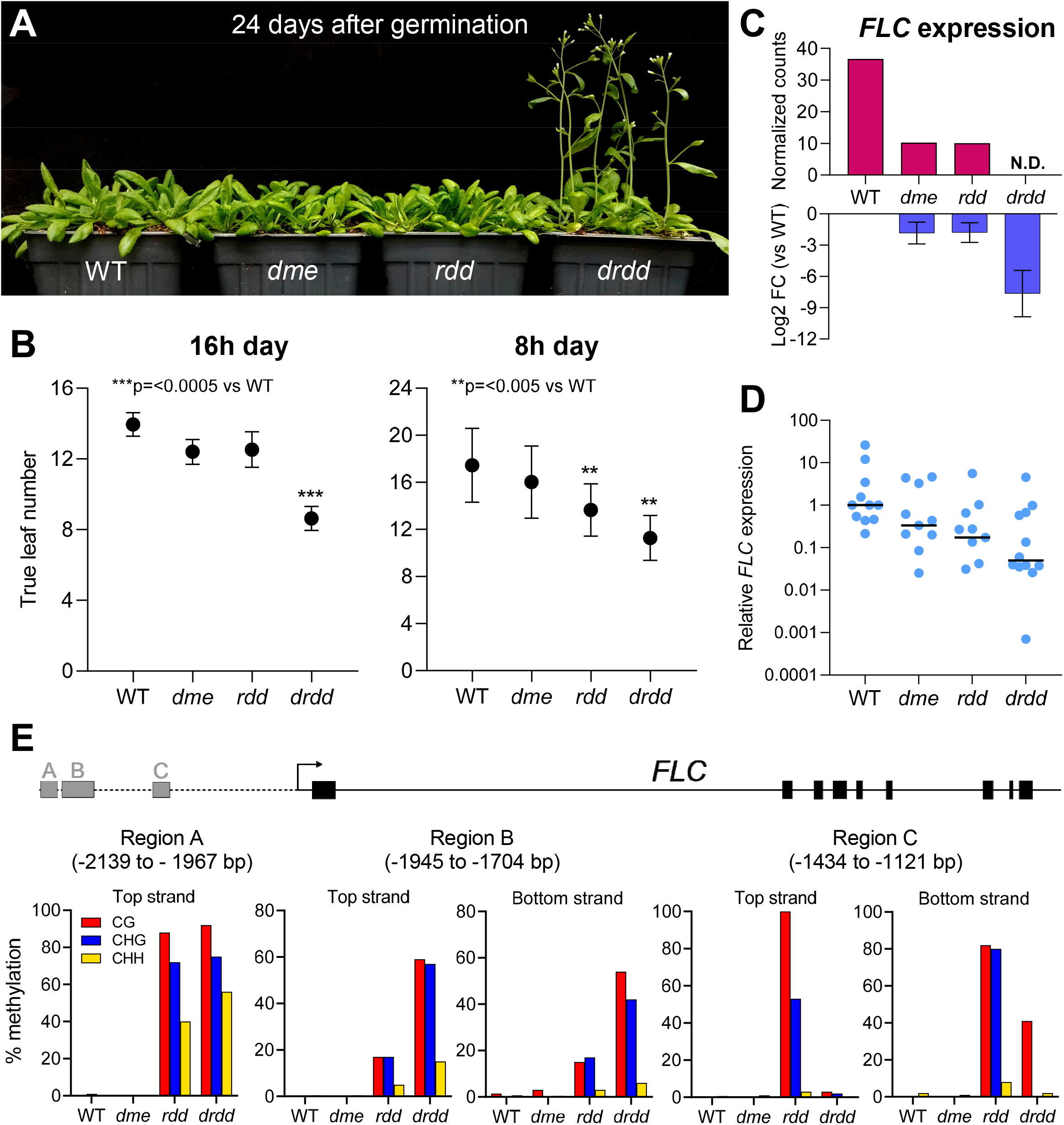
DRDD enzymes prolong flowering time and remove DNA methylation proximal to *FLOWERING LOCUS C*. A) Flowering phenotype of long-day grown WT, *dme, rdd* and *drdd* plants 24 days after germination. B) Flowering time of WT, *dme, rdd* and *drdd* plants grown in long (16 h) and short (8 h) day conditions. Flowering time is measured by the number of true leaves present at the initial time of bolting. Error bars represent standard deviation between biological replicates. C) *FLC* expression and fold-change compared to WT as measured by RNA-seq. Error bars represent standard error between biological replicates. D) Relative *FLC* expression measured by quantitative real-time PCR. Each point represents an individual biological replicate, horizontal lines denote the median value. E) Bisulfite-PCR analysis of methylation levels at three regions 5’ of *FLC*.

## Discussion

In this study we sought to elucidate the role of the DME family of 5-mC DNA glycosylases in somatic tissues of *Arabidopsis thaliana*. To test this, we generated somatic null mutants of *dme* as well as quadruple *drdd* mutants, abolishing the entire active demethylation family for the first time. Null *dme* mutants exhibited similar phenotypes, DNA methylation profiles, and transcriptional profiles to WT plants, strongly suggesting that DME does not perform a distinct individual function within leaf tissue. However, when combined with mutations to the other three members of the DME-family to generate *drdd* mutants, a number of previously unknown targets and functions for active demethylation in somatic tissues were brought to light.

### Active DNA demethylation targets a subset of gene bodies

Compared to WT, *drdd* mutants exhibited DNA methylation gains at 2,600 loci. We estimate the true number of DRDD targets to be even higher, as we employed stringent cutoffs for both read depth and the effect size of methylation changes. Approximately half of the DRDD targets gained methylation in all three sequence contexts, likely deposited by the RdDM pathway. These targets are similar in manner to previously described RDD targets – loci proximal to TEs or intergenic sequences that are targeted by RdDM, and exhibit an increase or “spreading” of DNA methylation in the absence of demethylase activity (Penterman et al., 2007; Lister et al., 2008; Tang et al., 2016). However, we also identified approximately 1,000 targets unlike those previously described in demethylation mutants, in which DNA methylation present in the CG-only context was removed. These regions are typically located within open chromatin, such as introns, exons, or untranslated regions and are referred to as gene body methylation (gbM). However, the sites we describe here are distinct from typical gbM sites, in which CG methylation at individual sites is close to 100% (Picard and Gehring, 2017). In WT plants, the CG-only regions we identified exhibit intermediate methylation levels, in which CG dinucleotides are methylated at 5-50% (Supp. Fig. 3). This low to intermediate methylation level implies that the CG sites are methylated in only a fraction of cells in the tissue. Although many of the CG-only target loci exhibit low methylation levels in WT leaves, they are more highly methylated in buds and embryos (Supp. Fig 4). Importantly, these loci are also highly methylated in *drdd* leaves, suggesting their methylation state is established by the active removal of methylation by DRDD enzymes in a subset of leaf cells. The function, if any, of gbM has been enigmatic. Globally, gbM is associated with moderately expressed genes (Zilberman et al., 2007), but some plant species have dispensed with this aspect of methylation patterning altogether (Bewick et al., 2016). Nevertheless, we observed 11% of down-regulated genes contained a CG-only DRDD target within their gene body, suggesting that methylation at these loci may indeed impact gene expression. Thus, we propose that active demethylation may act in a specific subset of cell types, cell cycle, or endoreduplication states in wild-type tissues, and that the intermediate methylation levels in WT plants are a consequence of heterogeneity in methylation state between cells. The absence of this heterogeneity within *drdd* mutants could lead to interesting future research on the establishment of methylation patterns within specific cell types.

The discovery of CG-only DRDD targets within gene bodies also raises questions about possible targeting mechanisms for DRDD demethylases. Previous work has proposed that active demethylation by ROS1 may be targeted by specific histone modifications associated with transcriptionally repressed intergenic sequences in Arabidopsis, such as H3K18ac and H3K27me3 (Tang et al., 2016; Qian et al., 2014). Our results are not fully consistent with this view, as we observed DRDD targets in transcriptionally active genes, and across a broad range of chromatin states (Fig. 2E) not dissimilar to the overall distribution of chromatin states in the Arabidopsis genome. It is possible that functional diversification between ROS1 and DME may contribute to such differences in their individual targeting, although the near-wild-type methylation profiles of single *dme* mutants suggests a high degree of redundancy in their activity and function.

### Active DNA demethylation impacts flowering time by targeting FLOWERING LOCUS C

*FLC* is a key regulator of the transition to flowering in *Arabidopsis*. The transcriptional silencing of *FLC* under prolonged exposure to cold mediates the vernalization response in some accessions via a Polycomb repressive complex switch (Costa and Dean, 2019). In addition, *FLC* also regulates the autonomous flowering pathway in early-flowering strains of Arabidopsis, such as Col-0 (Wu et al., 2020). *FLC* is expressed in the shoots of Col-0 plants in the initial weeks after germination, but expression decreases immediately prior to flowering (Pien et al., 2008; Choi et al., 2009). *FLC* has been established as a model locus for epigenetic and transcriptional control, and is beginning to be understood with a high degree of complexity (Wu et al., 2020). Despite extensive previous research, connections between *FLC* regulation and DNA methylation have been limited and indirect. While changes to *FLC* transcript abundance have been reported in some DNA methyltransferase and DNA methylation-binding mutants (Sheldon et al., 1999; Finnegan et al., 2005; Peng et al., 2006; Yaish et al., 2009), targeting of the *FLC* locus by either methylation or demethylation pathways has not previously been reported.

*FLC* is one of the most strongly down-regulated genes identified by RNA-seq in *drdd* compared to WT. This expression change was fully consistent with the clear early flowering phenotype of *drdd* mutants in both long and short day photoperiods. Unlike previous studies linking *FLC* expression to DNA methylation, we also observe a clear difference in methylation patterning proximal to the *FLC* locus that could feasibly explain the observed transcript abundance (Fig. 5E). It is possible that DNA methylation at this region could interfere with the binding of key regulators of *FLC* transcription. The FLC regulator FRIGIDA has been shown to form a super-complex with the H3K4 methyltransferase complex COMPASS-like at *FLC* (Li et al., 2018). Binding of this complex has been detected at a broad region encompassing 500 bp on either side of the *FLC* transcriptional start site. While this region does not directly overlap the sequences hypermethylated in *rdd* and *drdd* mutants, it is possible that demethylation of this region is required to establish the chromatin landscape necessary to facilitate binding of this super-complex.

As *FLC* appears to be targeted by DRDD, yet is completely unmethylated in wild-type plants, an interesting question is whether this intergenic region is co-targeted by methylation and demethylation pathways in wild-type leaves, resulting in a constant battle between the addition and removal of DNA methylation. Active DNA demethylation at this locus may maintain epigenetic homeostasis (Williams and Gehring, 2020) by preventing the over-reach of methylation-targeting pathways, or methylation of *FLC* might be important for modulating expression at some point in the life cycle. Small RNAs with homology to the 5’ intergenic region of *FLC* (Lee et al., 2012; Jeong et al., 2013; Erdmann et al., 2017) could drive the activity of the RdDM pathway, only for methylated cytosines to be subsequently removed by DRDD. Furthermore, *FLC* is an imprinted gene in the endosperm of some *Arabidopsis* accessions (Pignatta et al., 2014), a phenomenon that has been observed at other loci with methylated 5’ regulatory intergenic regions. Understanding why dynamic methylation and demethylation pathways appear to be engaged in a stalemate at this locus will be an exciting avenue for future research.

In summary, we propose that active demethylation in somatic tissues by DRDD plays an important role in maintaining epigenetic states that can influence transcriptional activity. This activity appears to be important in protecting the chromatin landscape at *FLC*, a locus at which a number of dynamic epigenetic mechanisms converge. Similar to reproductive development, active demethylation by DRDD may also be developmentally regulated, and may act as a mechanism to establish divergent epigenetic states between cell and tissue types.

## Methods

### *pAGL61::DME* transgene

The transgene to rescue DME expression in central cells was created by amplifying the promoter of *AGL61* (F primer: TCTAGAGGATCCAACCGATTTGACAA, R primer:

TGATCGCTAGCTCCTCCTTTTGTA), the full genomic coding sequence (introns included) of DME (F primer: ATGAATTCGAGGGCTGATCCG, R primer: TTAGGTTTTGTTGTTCTTCAATTTGCTC) and cloning both fragments into pENTR-TOPO-D via Gibson assembly (overhang sequences are not included in the primers above). The assembled *pAGL61:DME* construct was then transferred to the binary vector pMDC99 (Curtis and Grossniklaus, 2003) using LR clonase.

### Plant material

Triple homozygous mutant *rdd* plants were transformed with *pAGL61:DME* via floral dipping (Clough and Bent, 1998). Single-insertion transformants were selected and pollinated with *dme-2* (Choi et al., 2002) heterozygote mutant pollen to generate F_1_ progeny heterozygous for all four DRDD demethylase genes. These quadruple heterozygous plants were self-fertilized, and over two subsequent generations of segregation the following genotypes were isolated, each homozygous for the *pAGL61:DME* transgene: *dme, rdd* and *drdd* and DRDD WT segregants (to serve as a closely related WT control for downstream experiments). The selfed progeny of the initial plant of each genotype was used for all downstream experiments. To assess seed abortion, siliques were harvested after drying and seeds examined under a dissecting microscope. *rdr2* plants were homozygous for the *rdr2-1* allele.

### Flowering time assay

Plants were sown such that every row of the flats contained one plant of each genotype (WT, *dme, rdd*, and *drdd*) with the order iterating by one with each successive row. Flats were grown in a Conviron CMP6050 Control System at 22°C and 50% relative humidity, with 16 hours of 120 μMol light and 8 hours of darkness per day. Starting at two weeks of age, all plants were visually inspected three times per week. The number of rosette leaves was recorded once a bolt was visible. Populations were compared using a one-way ANOVA with post-hoc Tukey test. In addition, the experiment was performed in a growth chamber with identical conditions, except for a short-day growth cycle with 8 hours of light and 16 hours of darkness. The long day flowering time assay was repeated three times independently, and the short day assay was repeated twice, using >30 biological replicates for each experiment.

### Bisulfite sequencing

Four replicate samples of the fifth true leaf of 3-week old WT, *dme, rdd* and *drdd* individual plants were collected (16 plants total). Leaf samples were split along the midvein, with one half each used for DNA and RNA extractions. DNA was extracted using a CTAB protocol, and bisulfite conversion was performed on 200 ng DNA using an Invitrogen MethylCode bisulfite conversion kit. Bisulfite-sequencing libraries were then generated using a QIAGEN Qiaseq Methyl Library Kit, with 11 cycles of amplification and each individual replicate separately indexed. Samples were then sequenced using an Illumina HiSeq 2500 rapid run using 100 × 100 bp paired-end protocol at the Whitehead Institute Genome Technology Core. All 16 samples were multiplexed equally in two separate lanes, to avoid batch effects in sequencing. To improve genomic coverage for downstream analyses, the reads for every two replicates were pooled, creating two high-depth biological replicates for each genotype.

To sequence different tissue types, two WT Col-0 and *rdr2* replicates were sampled throughout each individual plant’s development. The fifth true leaf was collected on day 21; the second cauline leaf and closed flower buds from the primary inflorescence were collected on day 35; and a pool of 30 mature green embryos were collected on day 50. DNA extraction, bisulfite conversion, library preparation, and sequencing was conducted as above, with the following modifications: 1) embryo samples yielded less than 200ng of DNA, so reduced quantities of 103 ng and 143 ng were used for each replicate; 2) libraries were amplified with 6-10 cycles; 3) samples were equally multiplexed across three lanes and sequenced using an Illumina HiSeq2500 standard run with 100 x 100 bp paired-end reads at the Whitehead Institute Genome Technology Core.

### RNA-seq

RNA samples were isolated from 3-week old leaf samples (explained above) using a QIAGEN RNeasy plant mini kit. 400 ng total RNA was used to generate RNA-seq libraries using a QIAGEN QIAseq Stranded mRNA Select Kit, with 13 cycles of amplification. An additional round of purification using QIAseq beads was performed to remove adapter dimers. RNA-seq was performed on an Illumina HiSeq 2500 using a 50 bp single-end protocol at the Whitehead Institute Genome Technology Core. All samples were multiplexed equally in two separate lanes to avoid batch effects in sequencing.

### DNA methylation analysis

Prior to mapping, adapters were trimmed using Trim Galore (Babraham Bioinformatics), trimming 8 or 10 bp from the 5’ end of reads, and enforcing a 3’ end quality of >25%. Reads were mapped to the Araport11 genome using Bismarck 0.20.1 (Krueger and Andrews, 2011), allowing for 1 or 2 mismatches per read and removing PCR duplicates. Methylation values for each cytosine were calculated using the Bismark methylation extractor function. The efficiency of bisulfite conversion was verified by quantifying the percentage of methylation for reads mapped to the chloroplast. Bisulfite conversion rates were >99.7% across all samples. Before identifying differentially methylated regions (DMRs), SNPs homologous to the Ws-2 genome were identified as described (Picard and Gehring, 2017), and their zygosity was plotted across each chromosome (Supplemental Figure 1). This is because the *ros1* and *dml2* T-DNA insertions in *rdd* were originally introgressed from a Ws-2 background. In order to exclude genomic regions (and associated methylation differences) originating from the Ws-2 ecotype, two regions were excluded from all subsequent methylation analyses: chromosome 2 (8,802,496–15,397,296 bp) and chromosome 3 (677,340–5,117,803 bp).

To identify differentially methylated regions (DMRs), the genome was divided into 200 bp windows overlapping by 100 bp. Symmetrical cytosines within CG base pairs were combined to make a single averaged data point, as the two opposite-stranded cytosines within CG base pairs are not statistically independent. A “methylation score” was then calculated for each window based on the density of differentially methylated cytosines (DMCs) within each window. This methylation score was calculated as described (Williams and Gehring, 2017), with the added stringency that DMCs must be present in both biological replicates for each sample. This filter ensured that only regions adequately covered and differentially methylated in all samples could be identified as DMRs. The minimum methylation difference for each cytosine context was as follows: CG – 35%, CHG – 20%, CHH 15%. Cytosines with fewer than 5 reads coverage in each sample being compared were excluded from the analysis.

DRDD targets were identified by combining all windows hypermethylated in *drdd* compared to WT in any sequence context and merging together using bedtools merge. DRDD targets were split into two categories based on their methylation profiles. CG-only targets were identified based on the absence of non-CG methylation (both CHG and CHH <5%) in *drdd*, whereas multi-context targets were defined by possessing methylation in more than one sequence context (>5% CHG and/or CHH). The presence of DRDD target loci in each chromatin state of the genome was determined by intersecting DRDD targets with the chromatin states identified by (Sequeira-Mendes et al., 2014). Chromatin states 4 and 5 were combined to represent “Intergenic polycomb” and chromatin states 8 and 9 were combined to represent “heterochromatin”.

### Gene expression analysis

Prior to mapping, adapters were trimmed using Trim Galore (Babraham Bioinformatics), trimming 9 bp from the 5’ end of reads, and enforcing a 3’ end quality of >25%. Reads were mapped to the Araport11 genome using STAR (Dobin et al., 2013). Differentially expressed genes were identified by running htseq-count and DE-seq2 (Love et al., 2014), ensuring a minimum of 2-fold change in expression and a Benjamini-Hochberg corrected p-value <0.05. Proximity between differentially expressed genes and DMRs was calculated using BEDTools (closest). As a negative control, the proximity between differentially expressed genes and 10 sets of randomly selected 200 bp windows were calculated for each comparison. For example, proximity to 2,601 DRDD target DMRs was compared with proximity to 2,601 random windows for which hypermethylation could have been detected (e.g. <65% CG, <80% CHG and <85% CHH methylation in wild-type). Genes mapping to chromosomal regions with homology to Ws-2 were omitted from the differential expression and DMR proximity analyses shown in Figure 4. GO term analysis was performed using DAVID (Dennis et al., 2003).

### Locus-specific bisulfite-PCR

DNA was extracted using a CTAB protocol, and bisulfite conversion was performed on 200 ng DNA using an Invitrogen MethylCode bisulfite conversion kit. Bisulfite PCR was performed using a hot-start Taq polymerase (either Ex-Taq HS (Clontech) or DreamTaq HS (Thermo Scientific)) with an annealing temperature of 50 °C and an extension temperature of 68 °C. PCR products were purified with a MinElute Gel Extraction Kit (QIAGEN) and cloned into the pJET1.2/blunt vector (CloneJET PCR Cloning Kit; Thermo Scientific). Individual colonies (15-25 per locus) were either PCR screened with pJET F and R primers, or plasmids were extracted with a QIAprep Spin Plasmid Miniprep Kit (QIAGEN). PCR screen amplicons or plasmids were sequenced by Sanger sequencing using the PJET1-2F universal primer. Sequences were checked manually for quality, and vector and primer sequences were removed with SnapGene (V5.2.4). Sequences were aligned with Clustal Omega (Sievers et al., 2011) and methylation state was analyzed using CYMATE (Hetzl et al., 2007). The following primers were used for each locus: FLC region A (F: ATTTGGTYAGTAYTYAATTTTTGTGGTA, R: CCCCACATCAATCCAARTTCAA); FLC region B (top strand F: TTGAAYTTGGATTGATGTGGGG, top strand R: TTCCCTCCAAACCAATTTRARTTTATTT, bottom strand F: CCACTRTACTACTTACATTTTAACTAC, bottom strand R: AAAGTGTAYTTATAAGYATTAGGTTGTT); FLC region C (top strand F: GTGTYTTGYYAAATTAATAAAAAGGTG, top strand R: CCACACAAACATTTCACTAACACT, bottom strand F: CTCCAATARAAAARTTAATACCAATCAT, bottom strand R: GAAAAGAAGTGGGTGAAAYTGATTA); SPCH (F: TAGGAGGAGTTGTGGAGTAYATAAG, R: CAARCTCATTRATCACACTACTCTCAT); SOC3 (F: GAAAAGAAGATGATGTTGTTYAAYAAGT, R: CCCTCCTTCATAARCTCCATTATC).

### Quantitative real-time PCR

RNA was isolated from the 5th leaf of 21-day old plants using TRIzol Reagent (Invitrogen) according to manufacturer’s instructions. Genomic DNA was removed by treatment with Amplification-grade DNase I (Invitrogen). cDNA was prepared from 500-750ng RNA (standardized within each batch) with Superscript II Reverse Transcriptase (Invitrogen) according to the manufacturer’s instructions, with polyadenylated transcripts selected for through use of an oligo-dT primer. qPCR was performed on a StepONE Plus Real-Time PCR system with Fast SYBR-Green PCR master mix (Applied Biosystems). FLC primer sequences were as previously described (Csorba et al., 2014): FLC_F: AGCCAAGAAGACCGAACTCA, and FLC_R: TTTGTCCAGCAGGTGACATC. Reactions were normalized to reference gene AT1G58050 (Czechowski et al., 2005). All qPCR reactions were performed with technical triplicates. Cycling conditions were as follows: 95°C for 20s followed by 40 cycles of 95°C for 3s and 60°C for 30s. Relative fold change in expression was determined using the ΔΔCt method (Livak and Schmittgen, 2001).

### Accession Numbers

All high-throughput sequencing data is deposited in NCBI GEO under accession GSEXXXX.

## Supplemental Material

**Supplemental Figure 1: Zygosity of Ws-2 SNPs in *rdd* and *drdd* mutants**

**Supplemental Figure 2: DNA methylation differences between tissue types**

**Supplemental Figure 3: Tissue-specific methylation levels of DRDD targets in WT and *rdr2* mutants**

**Supplemental Figure 4: Significantly enriched GO terms in *drdd* versus WT**

**Supplemental Figure 5: Bisulfite PCR validation of CG-only DRDD targets**

**Supplemental Figure 6: Additional flowering time assay replicates**

**Supplemental Dataset 1: List of DRDD target regions**

**Supplemental Dataset 2: Differentially expressed genes identified by DESeq2**

## Acknowledgements

We are grateful to Enrico Calvanese for performing bisulfite PCR validation experiments and Maria José Aranzana for performing a flowering time replicate. Research in this publication was supported by the National Institute of General Medical Sciences of the National Institutes of Health under award R01GM112851 to M.G.

## Author Contributions

BW and MG conceived the study. BW, LB, and DP designed and performed experiments and data analysis. BW and MG wrote the paper.

