## Supplementary Figures for "Somatic DNA demethylation generates tissue-specific methylation states and impacts flowering time"

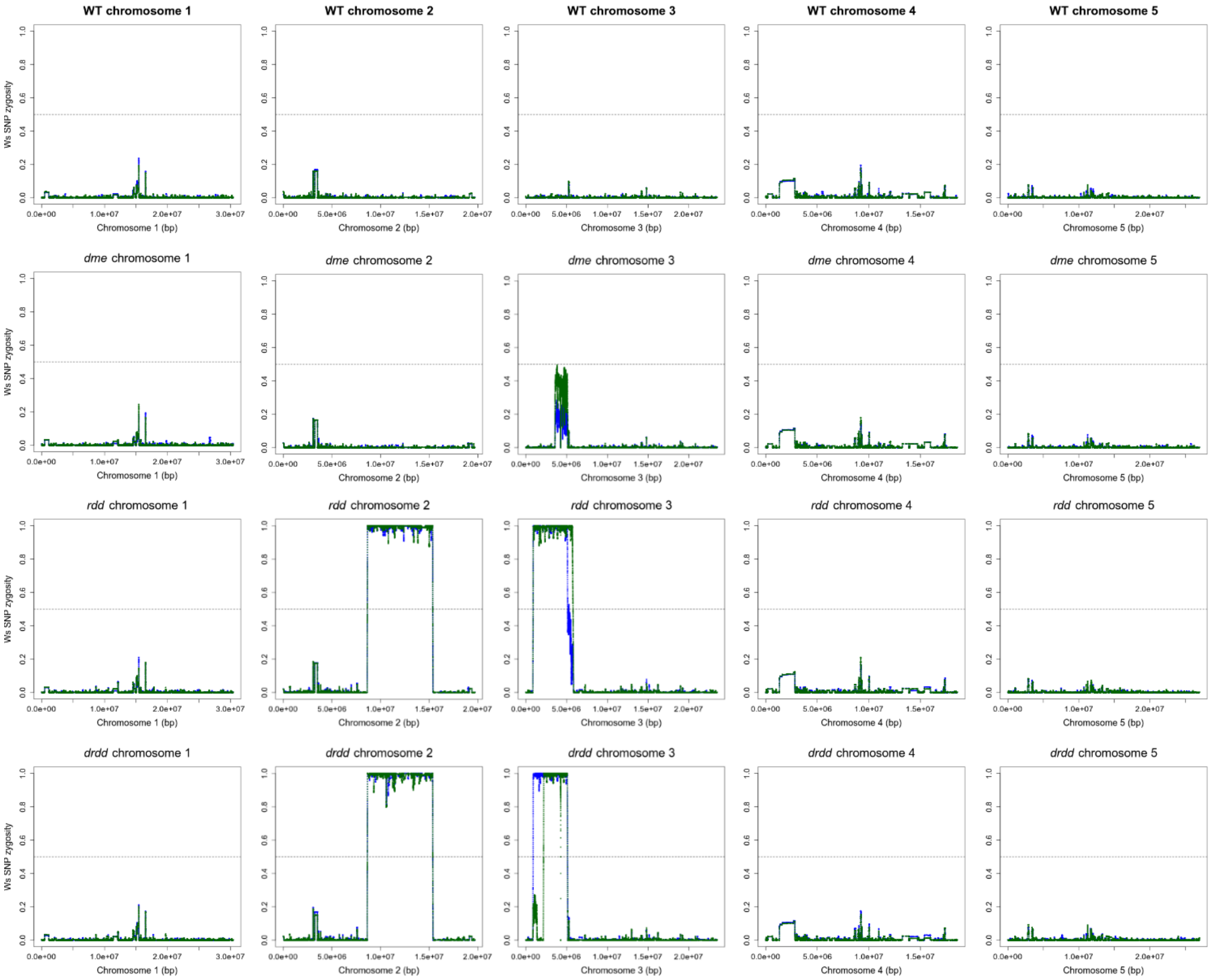


**Supplemental Figure 1: Zygosity of Ws-2 SNPs in *rdd* and *drdd* mutants**

Zygosity of Ws-2 SNPs in whole genome bisulfite-seq data is plotted for each chromosome in WT, *dme*, *rdd* and *drdd* mutants. Green and blue points represent each of two biological replicates.

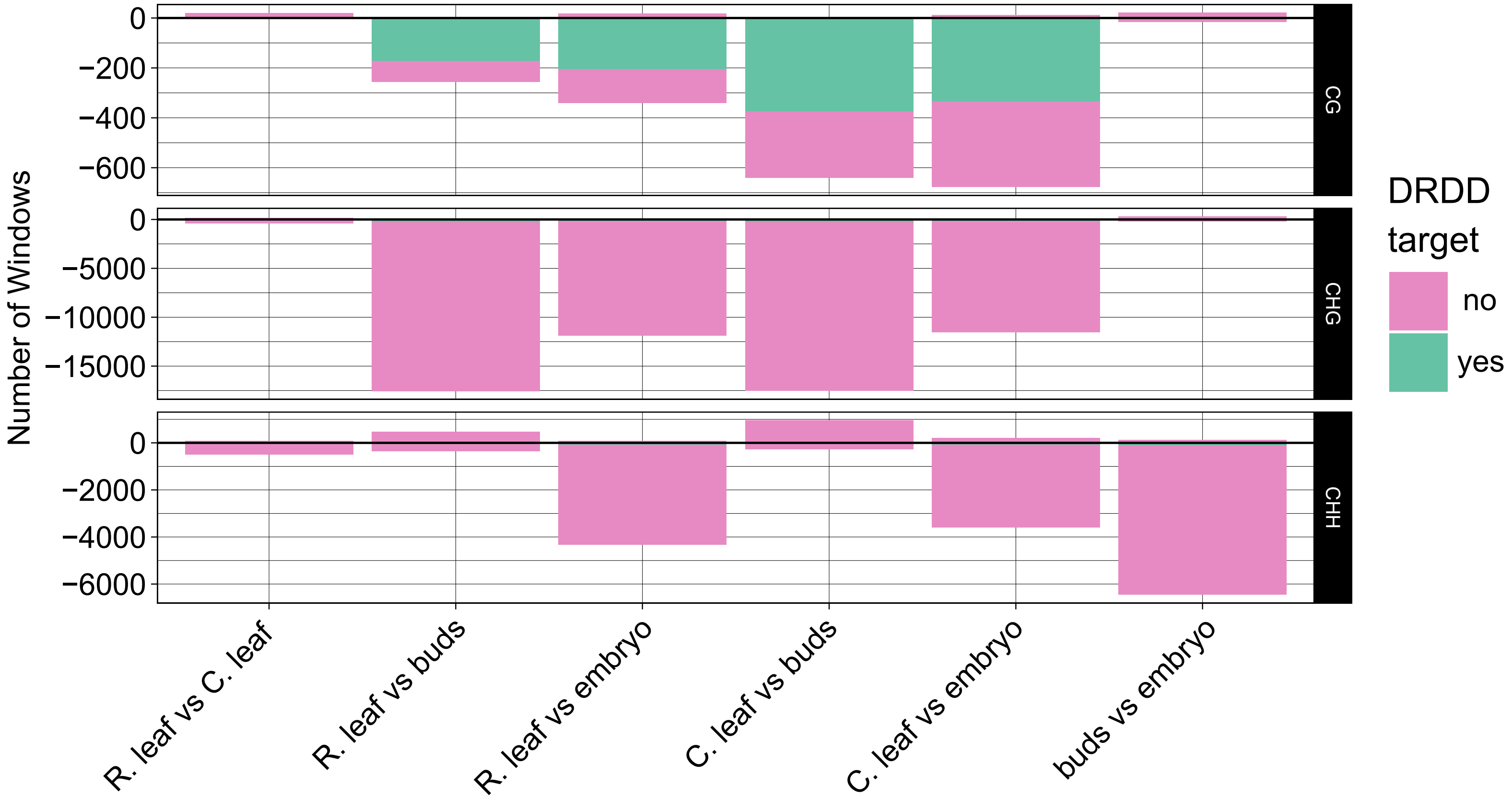


**Supplementary Figure 2: DNA methylation differences between tissue types**

The number of hypermethylated windows is plotted above zero, whereas the number of hypomethylated windows is plotted below zero. The proportion of windows overlapping with DRDD target loci is plotted in green. R. leaf = rosette leaf, C. leaf = cauline leaf

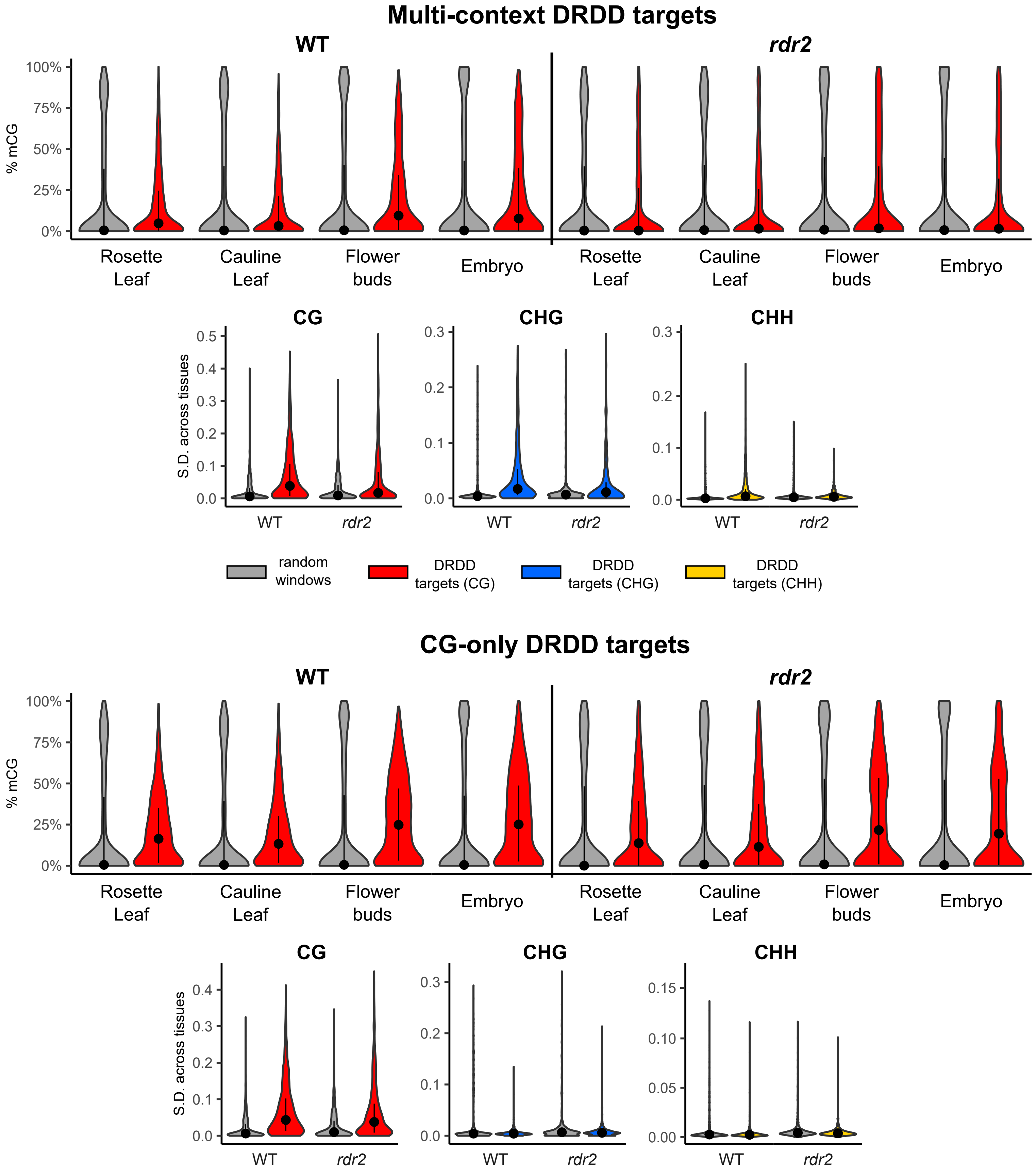


**Supplemental Figure 3: Methylation levels of DRDD targets across tissues in WT and *rdr2* mutants**
Violin plots showing CG methylation levels at multi-context and CG-only DRDD targets in four different tissues of WT and *rdr2*. DRDD targets are compared against 2,601 randomly selected genomic windows. Violin plots of the standard deviation of CG, CHG and CHH methylation levels across all four tissues are also shown below. Within violin plots, black dots represent the median value and black lines denote the interquartile range.

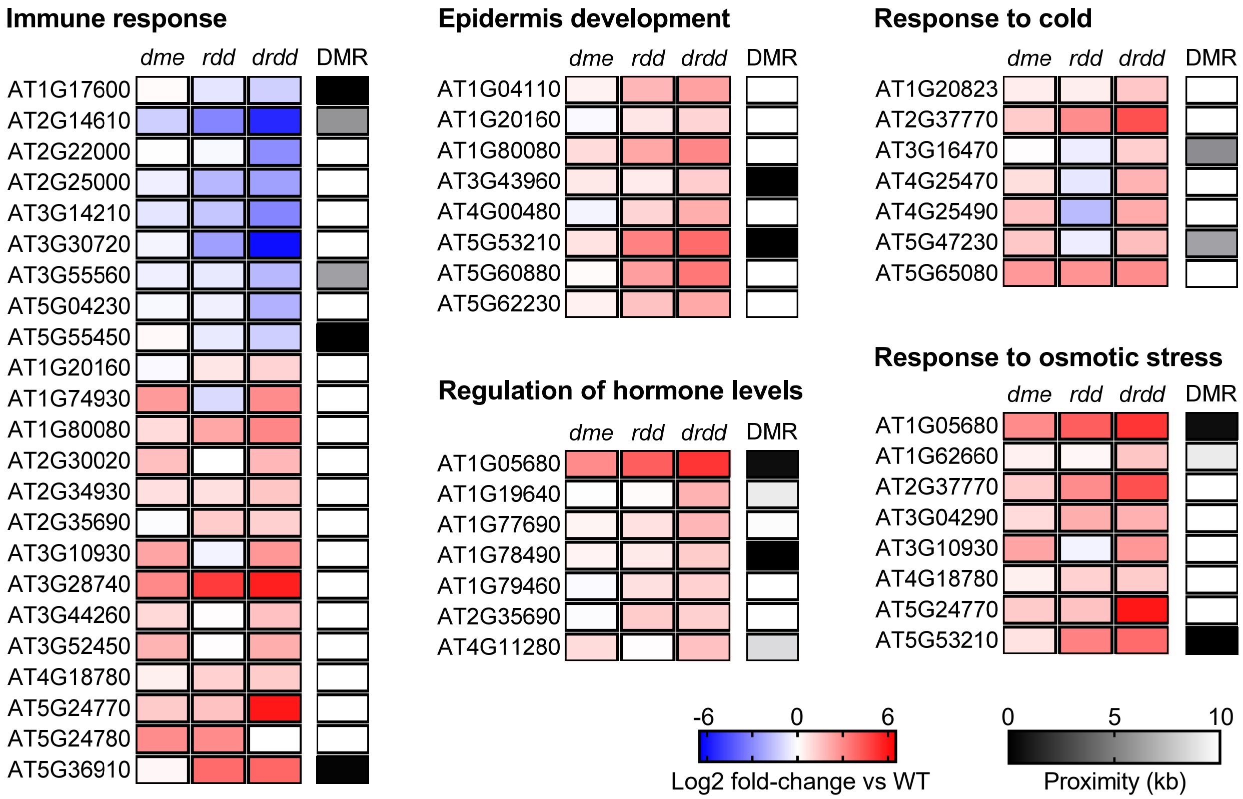


**Supplemental Figure 4: Significantly enriched GO terms in *drdd* versus WT**
Log2 fold-change gene expression differences are plotted for differentially expressed genes within several functional categories enriched within *drdd* versus WT. Proximity to the nearest differentially methylated region (DMR) hypermethylated in *drdd* versus WT is shown adjacent to expression differences.

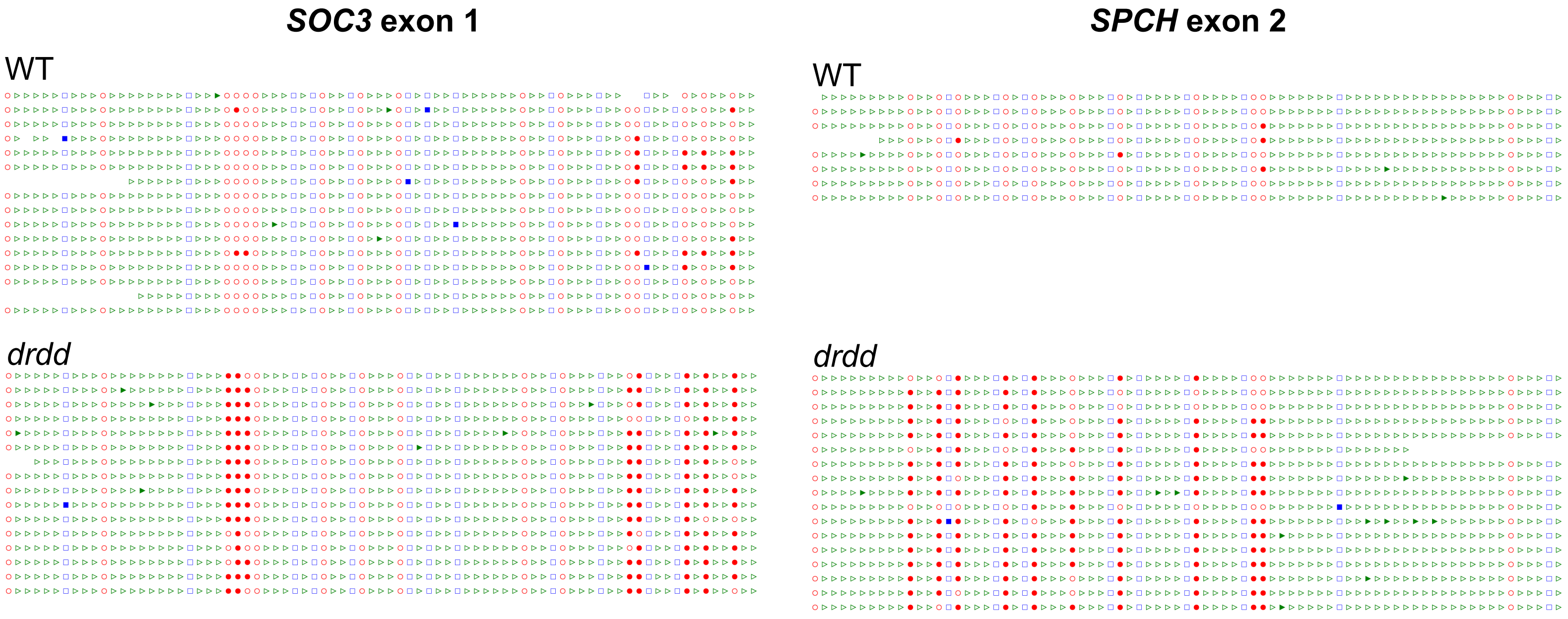


**Supplemental Figure 5: Bisulfite-PCR validations of CG-only DRDD targets**

Bisulfite PCR analysis of two CG-only DRDD targets, *SOC1* and *SPCH*. Filled and unfilled red circles denote methylated and unmethylated CGs respectively. Blue squares represent CHG sites and green triangles represent CHH sites. Each row represents one independent sequenced clone from each bisulfite PCR

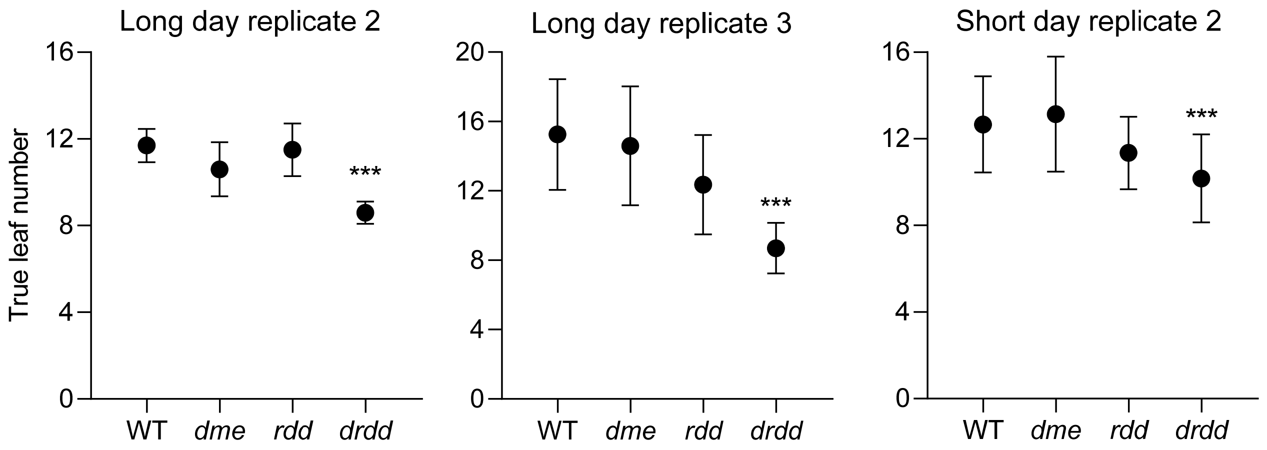


**Supplemental Figure 6: Additional flowering time assay replicates**
Additional replicate experiments are shown for both long day (16 h light) and short day (8 h light) flowering time assays. At least 30 plants were assayed for each genotype in each replicate experiment. ***p=<0.0005
